# Systematic re-annotation of 191 genes associated with early-onset epilepsy unmasks *de novo* variants linked to Dravet syndrome in novel *SCN1A* exons

**DOI:** 10.1101/648576

**Authors:** Charles A. Steward, Jolien Roovers, Marie-Marthe Suner, Jose M. Gonzalez, Barbara Uszczynska-Ratajczak, Dmitri Pervouchine, Stephen Fitzgerald, Margarida Viola, Hannah Stamberger, Fadi F. Hamdan, Berten Ceulemans, Patricia Leroy, Caroline Nava, Anne Lepine, Electra Tapanari, Don Keiller, Stephen Abbs, Alba Sanchis-Juan, Detelina Grozeva, Anthony S. Rogers, James Wright, Jyoti Choudhary, Mark Diekhans, Roderic Guigó, Robert Petryszak, Berge A. Minassian, Gianpiero Cavalleri, Dimitrios Vitsios, Slavé Petrovski, Jennifer Harrow, Paul Flicek, F. Lucy Raymond, Nicholas J. Lench, Peter De Jonghe, Jonathan M. Mudge, Sarah Weckhuysen, Sanjay M. Sisodiya, Adam Frankish

## Abstract

The early infantile epileptic encephalopathies (EIEE) are a group of rare, severe neurodevelopmental disorders, where even the most thorough sequencing studies leave 60-65% of patients without a molecular diagnosis. Here, we explore the incompleteness of transcript models used for exome and genome analysis as one potential explanation for lack of current diagnoses. Therefore, we have updated the GENCODE gene annotation for 191 epilepsy-associated genes, using human brain-derived transcriptomic libraries and other data to build 3,550 novel putative transcript models. The extended transcriptional footprint of these genes allowed for 294 intronic or intergenic variants, found in human mutation databases, to be reclassified as exonic, while a further 70 intronic variants were reclassified as splice-site proximal. Using *SCN1A* as a case study due to its close phenotype/genotype correlation with Dravet syndrome, we screened 122 people with Dravet syndrome, or a similar phenotype, with a panel of novel exon sequences representing eight established genes and identified two *de novo SCN1A* variants that now, through improved gene annotation can be ascribed to residing among novel exons. These two (from 122 screened patients, 1.6%) new molecular diagnoses carry significant clinical implications. Furthermore, we identified a previously-classified *SCN1A* intronic Dravet-associated variant that now lies within a deeply conserved novel exon. Our findings illustrate the potential gains of thorough gene annotation in improving diagnostic yields for genetic disorders. We would expect to find new molecular diagnoses in our 191 genes that were originally suspected by clinicians for patients, with a negative diagnosis.

## Introduction

The Early Infantile Epileptic Encephalopathies (EIEE) are a group of rare neurodevelopmental disorders, characterized by developmental delay or regression, electroencephalographic abnormalities, early-onset seizures that are often intractable and, in some cases, early death^1,2^. Large-scale international research efforts such as Epi25 <http://epi-25.org/>, the Deciphering Developmental Disorders (DDD) study^3^ and the UK 100,000 Genomes Project^4^ are now concentrating on diagnosing patients and identifying genes involved in rare disorders including EIEE, using chromosomal microarrays, whole exome sequencing (WES) and whole genome sequencing (WGS). However, while numerous novel genes associated with EIEE are being uncovered, between 60-65% of patients remain without a molecular diagnosis^5,6^.

Improvements in diagnostics can, in part, be achieved through improvements in gene annotation. Currently, most GENCODE (i.e. Ensembl)^7^ and RefSeq^8^ gene annotations are based on cDNA and EST libraries produced alongside the initial experimental phase of the human genome project. Such datasets were limited by a lack of human-specific sequence and relied on computer algorithms that used cross-species homology to identify, for example, putative open reading frames^9-11^ However, large datasets produced by more recent RNA-seq and long-read sequencing-based projects from human tissue remain largely unincorporated. Such datasets have the potential to add novel exons *via* transcript models to existing gene annotations, and these novel features can provide new insights into genetic disease. For example, ‘expanded’ exomes offer the potential to capture additional disease-linked mutations beyond the reach of previous studies. Additional sequences can be added to existing exome ‘panels’ used for diagnostics in the clinic and used to select novel regions for resequencing in patients without a molecular genetic diagnosis. Indeed, this has already been demonstrated for *DLG2*, where newly-identified exons were observed to be deleted in patients with neurodevelopmental disorders^12^. Furthermore, the resequencing of novel annotated regions identified previously-missed pathogenic variants linked to epilepsy in *SCN8A*^*13*^. Meanwhile, additional transcript features can also be used to reappraise existing variation datasets; providing, for example, new mechanistic explanations for known disease-associated variants, whilst also allowing for the reconsideration of variants from whole-genome studies that had previously been de-prioritised due to an apparent lack of transcription.

However, although modern transcriptomics datasets have the potential to add an *enormous* amount of novel transcribed features to gene annotation catalogues, the exact proportion of this ‘transcriptional complexity’ that is linked to gene function, and therefore to phenotype, remains hard to fathom. Thus, if a portion of observed transcription events for a given gene lack functional relevance, then expanded gene annotations could unknowingly be a source of misleading variant interpretations. In practice, transcript functionality is confirmed in the laboratory, although important insights can be gained from bioinformatics. Evolutionary conservation has long been regarded as a strong proxy for functionality, for example in the observation that coding sequences (CDS) have been subjected to purifying selection, or that splice sites are constrained and consistently transcribed in multiple species^14^. As such, conservation metrics are commonly used in transcript interpretation and thus variant prioritisation. However, it is becoming increasingly clear that variants linked to poorly conserved transcript features can also be drivers of genetic disease^11^. In particular, it is now well established that many genes utilise alternative splicing to reduce their translational output, redirecting transcription into non-coding pathways *via* the incorporation of ‘poison exons’^15^ While at least some poison exons are ancient, the mode and tempo of regulatory splicing evolution in general remains poorly understood. Alternatively, several recent reports have demonstrated that gene output can be compromised by mutations that improve the splicing efficacy of poorly conserved transcription events at the expense of the ‘canonical’ mRNA, i.e. according to a ‘gain of function’ model^16^

Here, we explore the potential of expanded gene annotation to improve the diagnostics of epilepsy, utilising a manual annotation workflow that is initially agnostic with regards to the potential functionality of the novel transcript sequences. Overall, this work has added 3,550 GENCODE transcript models to 191 genes associated with epilepsy *via* the utilisation of publicly available short and long read transcriptomics datasets. We subsequently create an expanded exome panel for eight genes associated with Dravet syndrome (an EIEE subtype), incorporating 125 newly-identified transcript regions, and, after resequencing 122 patients with a clinical diagnosis of Dravet syndrome or a similar phenotype, we discover two *de novo* variants within novel *SCN1A* exonic sequences. Both variants are found within presumptive poison exons that exhibit poor evolutionary conservation. In contrast, we also demonstrate that a third Dravet-associated variant, previously considered to be intronic and of unknown significance, is present in an alternative *SCN1A* poison exon that has deep conservation.

Although further work is required to understand the biological implications of the transcriptional complexity associated within *SCN1A* and the larger set of 191 genes, our findings relating to Dravet syndrome show that from the current bioinformatics perspective, uncertainties regarding transcript functionality should not necessarily be a barrier to the attribution of these transcripts in disease genetics.

## Results

### Manual re-annotation substantially increases the number of novel transcripts

We manually re-annotated 191 genes associated with epilepsy as part of the GENCODE project, primarily using publicly available long-read transcriptional datasets, including brain-derived SLR-Seq and PacBio Iso-Seq reads (see Methods). In total, 3550 novel alternatively spliced transcripts were annotated across the 191 genes, increasing the number of transcripts for these genes from 1807 to 5357. All novel transcripts are included in GENCODE v28; contemporary RefSeq annotation contains 1397 NM and XM transcripts. The majority of added transcripts contain either completely novel exons (37%) or utilize newly-identified alternative splice sites within existing exons (21%). In total, more than 674 kb of additional exonic sequence was added across the 191 genes.

Next, given the epilepsy context of this study, we further characterised the novel transcript regions using existing RNA-Seq data from pre-frontal cortex in 36 individuals across 6 life stages^17^. Novel introns are generally less well supported than pre-existing introns (Figure 1), and just 19% of introns between novel exons annotated as coding are covered by at least 10 RNA-Seq reads. However, while most detectable novel introns displayed a broadly ubiquitous temporal expression pattern, a subset of 208 (8%) showed a fivefold enrichment in expression in fetal versus infant pre-frontal cortex samples, with 101 showing more than fivefold higher expression than all other samples combined. This enrichment may highlight a subset of transcripts of particular functional importance. However, we recognise that the utility of tissue-specific expression patterns to infer functionality remains debatable^18,19^.

**Figure 1:**
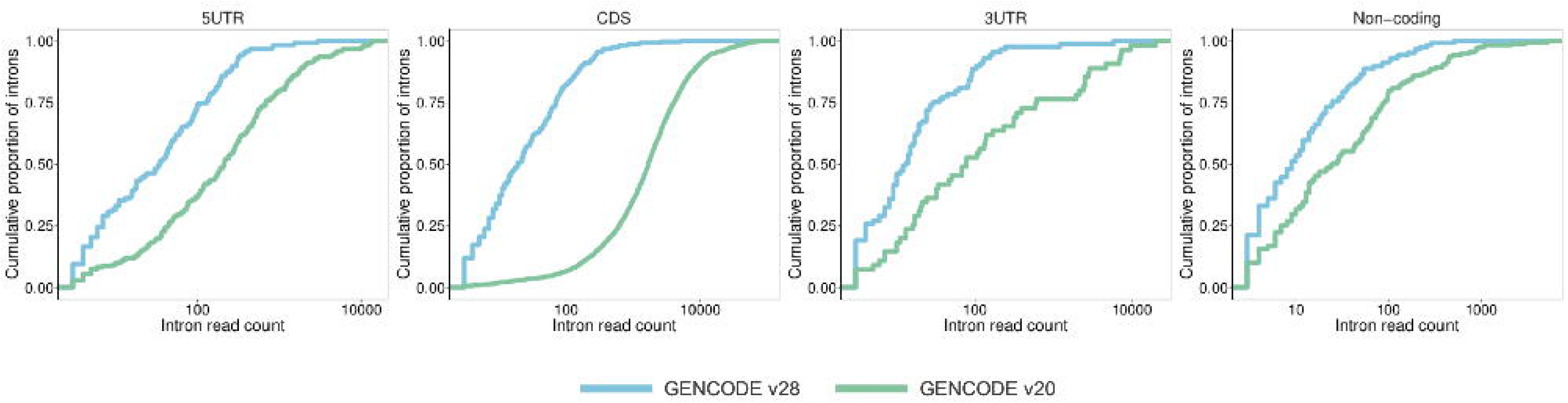
Expression of novel transcripts. Cumulative distribution curves for the number of intron supporting reads in pre-existing (GENCODE v20) versus updated (GENCODE v28) annotation. Distribution curves for overall transcripts, CDS, 5’ and 3’ UTR are given. The x-axis is in log10 scale.

Evolutionary conservation is a commonly used metric of functional potential^20^. Firstly, we mapped all human annotation for these 191 genes to the mouse reference genome using TransMap^21^. 40 % of novel introns mapped to the mouse genome with the preservation of canonical splice sites, compared to 87% of introns in the pre-existing annotation, indicating that the newly annotated introns show generally lower conservation. We next examined whether there was a correlation between general evolutionary conservation and the decision to annotate a novel transcription event as coding. We found that 18% of the additional exonic coverage in the novel annotation overlaps with PhastCons elements^22^, i.e. regions of the genome exhibiting detectable sequence conservation, compared to 47% of pre-existing exons. When considering only the 42kb of novel sequence annotated as coding, this proportion rises to 23%. However, just 6% of all novel CDS exhibits the characteristic pattern of protein sequence evolution, as judged by an examination of overlap with regions of positive PhyloCSF score^23^, compared to 91% of pre-existing coding exons. Although this may suggest that our annotation has significantly overestimated the proportion of the novel transcribed sequence that is translated, it could be that certain newly identified CDS regions have arisen in the primate lineage following the divergence from the rodent clade.

In summary, while expression, conservation and proteomics-based approaches do not provide vigorous support for the existence of *widespread* functionality across the novel transcribed regions, they do suggest that the reannotation of 191 genes has added a significant subset of models with conserved biological roles.

### Updated annotation identifies putative clinically-relevant variants

Given these considerations, we decided to investigate the clinical impact of the novel annotations without initial regard to their expression, conservation, or proteomics metrics. Firstly, we compared the overlap of pre-existing and updated annotation with five public variation datasets (dbSNP, ClinVar, ESP, HGMD-PUBLIC and PhenCode). We note that human mutation catalogues are currently biased towards ‘known’ gene sequences in terms of content, i.e. disease-associated mutations are less likely to be found in regions that are less well studied (or have been resequenced less often). Nonetheless, the updated annotation reclassifies 294 variants as exonic, and demonstrates that a further 70 intronic variants are found within 8bp of newly identified splice-sites (Table 1).

**Table 1.** The updated annotation reclassifies the impact of known variants in five different variant databases (dbSNP, ClinVar, ESP, HGMD-PUBLIC and PhenCode). Exon coverage is shown in grey shading on the left, while the intronic flank coverage for regions within 8 bp of the splice site is shown on the right hand side with no shading. Note that the numbers of variants overlapping the GENCODE v20 annotation and the novel annotation do not add up to the variant count in GENCODE v28 because not all pre-existing variants remain in the updated annotation. This is because the annotation update has also reclassified some previously exonic sequence as intronic.

|  | GENCODE v20 | GENCODE v28 | Novel | GENCODE v20 | GENCODE v28 | Novel |
| --- | --- | --- | --- | --- | --- | --- |
| ClinVar | 2200 | 2281 | 103 | 154 | 144 | 14 |
| ESP | 190 | 231 | 41 | 26 | 25 | 2 |
| HGMD-PUBLIC | 6897 | 7025 | 143 | 704 | 714 | 53 |
| PhenCode | 477 | 484 | 7 | 26 | 26 | 1 |
| dbSNP | 152693 | 220130 | 68892 | 8708 | 9699 | 2132 |
| total | 162457 | 230151 | 69186 | 9618 | 10608 | 2202 |
| total unique | 157492 | 224814 | 68795 | 9122 | 10104 | 2156 |

Secondly, we focussed on a specific form of epilepsy linked to a limited set of genes. Dravet syndrome is one of the best described and genetically most homogeneous EIEE syndromes^24^. More than 80% of Dravet syndrome cases are attributable to variants in *SCN1A*^*25*^ (OMIM ID: 607208) and about 700 pathogenic CDS variants have been reported^26^. Given the clear link between variants in *SCN1A* and Dravet syndrome, any un-annotated exonic sequence is of potential clinical interest.

The updated *SCN1A* annotation identified 10 novel exons and five novel shifted splice junctions (Figure 2; Table 2), increasing the genomic footprint of *SCN1A* transcription by approximately 3kb.

**Table 2.**
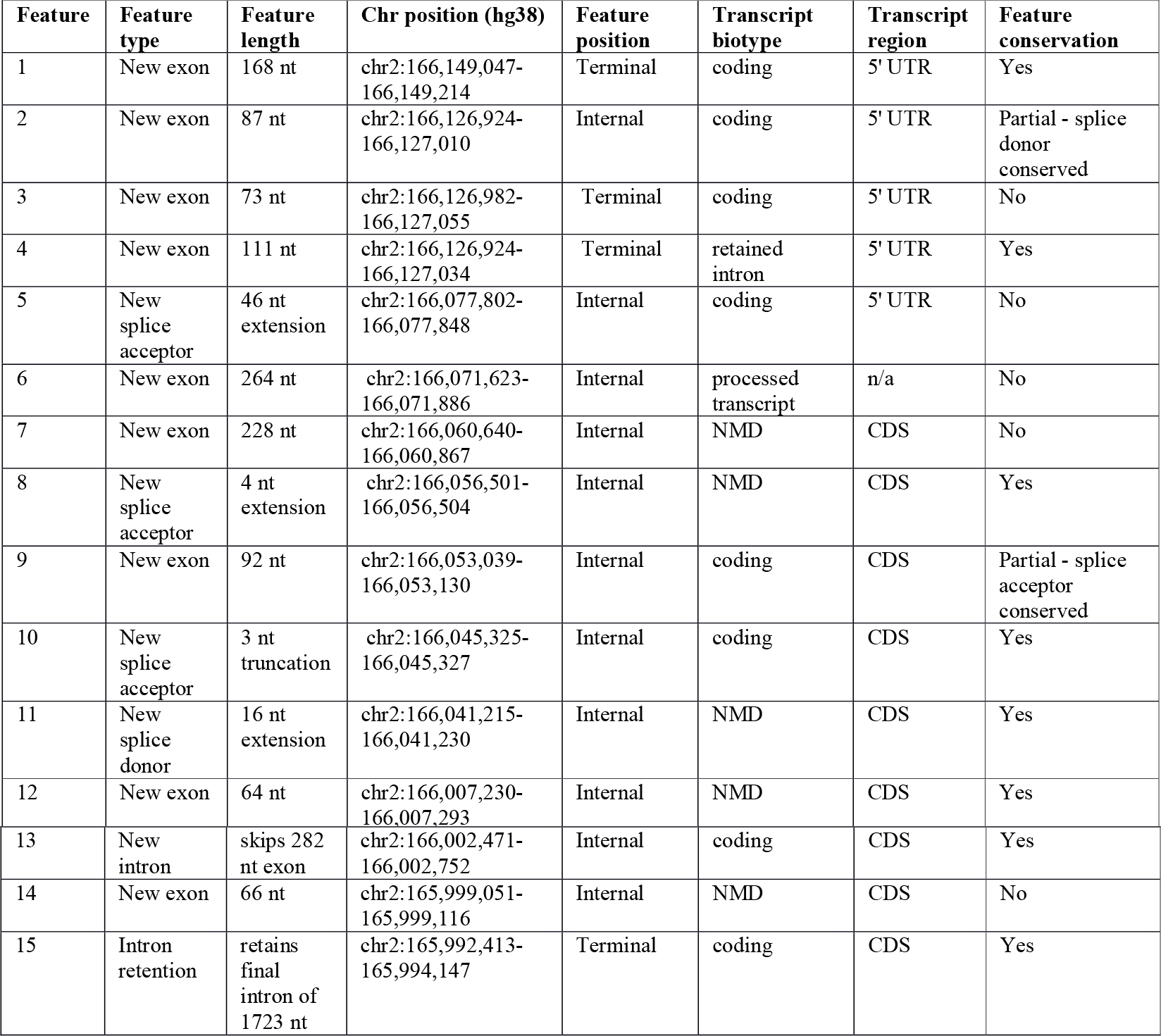
List of all novel features identified within *SCN1A*. A single Ensembl transcript model is listed for all novel features; certain features are also present in models. ‘Biotype’ details the functional effect of the feature as inferred by manual annotation. The alternative final exon within model ENST00000642141 was annotated as ‘non-coding’ due to the absence of polyadenylation data as per GENCODE guidelines; the functional status of this model is in reality unknown. Feature conservation describes the annotation and structurally identical feature in the mouse ortholog *Scn1a*. Chromosome coordinates are for the GRCh38 assembly. NMD is the abbreviation for nonsense mediated decay

**Figure 2:**
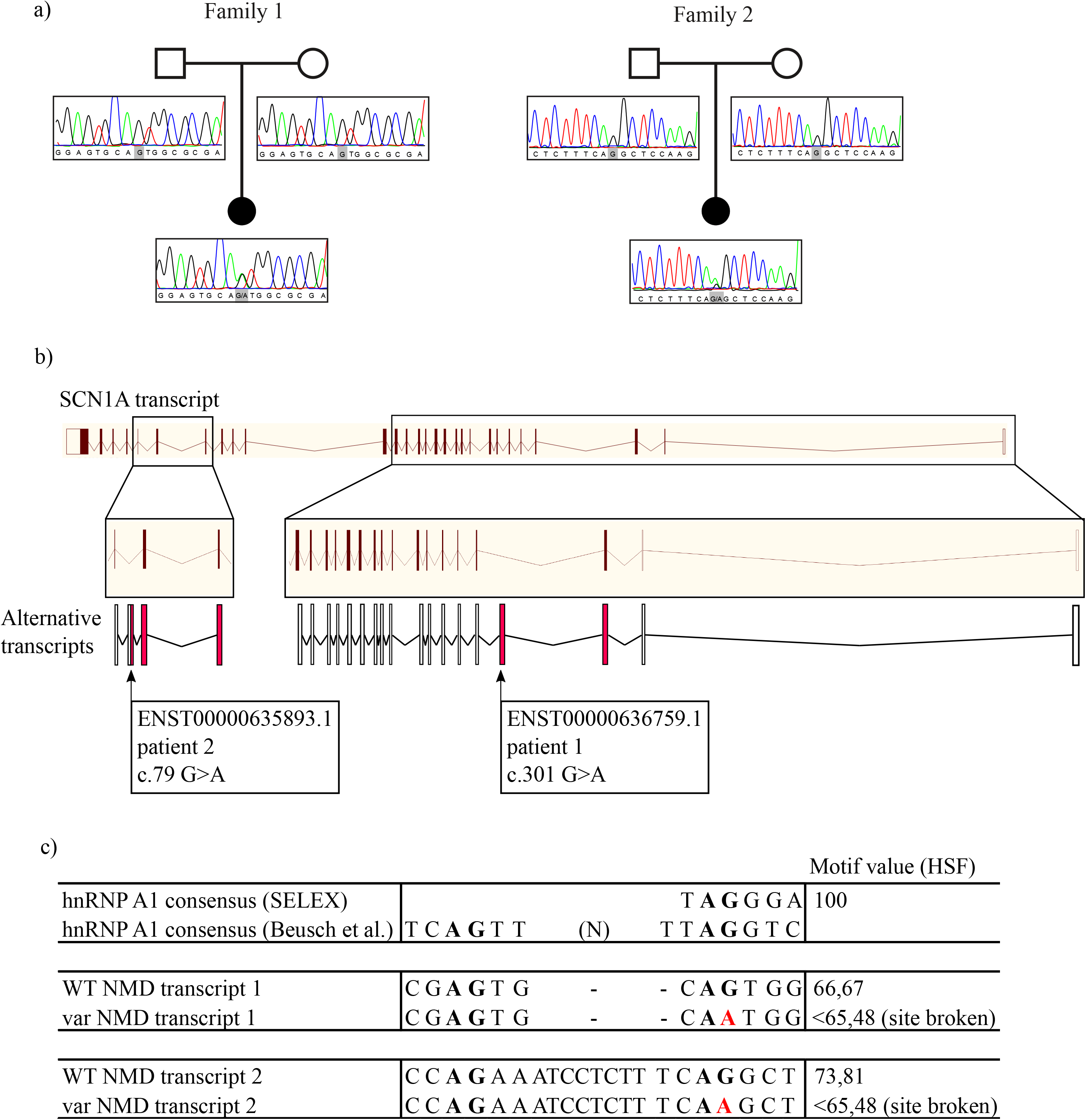
The updated *SCN1A* annotation identified 10 novel exons and five novel shifted splice junctions, increasing the genomic footprint of *SCN1A* transcription by approximately 3kb. All features are described with respect to existing Ensembl model ENST00000303395 and numbered according to the scheme used in Table 2. For clarity, the novel features are shown as truncated models containing only the exons of specific interest (and certain features are present on multiple transcript models in the complete gene annotation). UTR sequences are shown in grey, coding or NMD regions in black. Features [1] and [2] represent previously unreported 5’ UTR sequences that have conservation and equivalent expression in mouse and chicken. Features [7] and [14] are newly identified cassette exons predicted to invoke NMD and contain the *de novo* mutations identified in the study within patients 1 and 2 respectively. Feature 9 is cassette exon that is an ancient duplication of coding exon 5, to which it is transcribed in a mutually exclusive manner; the clinical significance of this exon has been previously demonstrated by Tate *et al*. Feature 12 is a cassette exon predicted to invoke NMD. Intron and exon sizes are to approximate scale. Additional transcript models have been omitted for clarity. (See Supplemental Data 1 for an illustration of all splicing features in table 2 for *SCN1A* in the Genoverse browser <https://wtsi-web.github.io/Genoverse/>)

At least two of these new annotations have demonstrably added biologically relevant sequences to the GENCODE catalogue. Firstly, we found that one novel exon had in fact already been reported in the human *SCN1A* literature, being described as alternatively spliced with respect to canonical exon 5 in a mutually exclusive manner^27^ (feature 9; Table 2, Figure 2). The inclusion of this alternative exon is known to generate *SCN1A* isoforms that differ in their expression pattern (with the novel exon preferentially expressed in neonatal brain, as confirmed by our analysis) and sensitivity to the antiepileptic medications phenytoin and lamotrigine. Nonetheless, it appears that this exon had not previously been included in any gene annotation catalogues and we therefore presume it has also been absent from *SCN1A* exome panels. Secondly, a novel poison exon (feature 12) was also missing from GENCODE, despite the fact that the orthologous exon in rat has been experimentally characterised^28^ (feature 12; Table 2, Figure 2). We report that this exon incorporates ClinVar variant RCV000209951, an *SCN1A de novo* variant found in a patient with Dravet syndrome. The variant had been annotated as intronic by ClinVar, and thus of unknown significance. Genome alignments support the conservation of this exon across vertebrate species. This variant has since been independently characterised as a gain of function mutation, promoting inclusion of the poison exon and leading to reduction in the amount of functional SCN1A protein^29^.

Two additional regions added to GENCODE as part of this study exhibit strong markers for functionality at the transcript level. The first is an alternative first exon consisting of 5’ UTR sequence, found approximately 21kb upstream of the previously recognised 5’ end of the *SCN1A* gene (feature 1; Table 2, Figure 2). The biological validity of this exon is underpinned by strong transcriptomics support in multiple data sources, and the fact that it is conserved in both mammalian and avian genomes, with brain-specific transcriptomics support in mouse and chicken. The second region represents additional 5’ UTR sequence, this time found as a cassette exon in the first intron following the recognised *SCN1A* first exon (feature 2; Table 2, Figure 2). Splice site conservation is observed across to avian genomes (although with a 3bp acceptor site shift in certain lineages), with brain-specific RNAseq support in chicken (not shown). Neither of these two novel regions overlaps with any known disease-linked variants. However, as previously stated, it should be noted that these regions have apparently been hitherto unstudied in a clinical context.

To further investigate the value of our novel annotations in epilepsy diagnostics, we screened a cohort of 122 patients with Dravet syndrome or a clinically similar phenotype, for the newly added regions of eight genes with known associations to this disorder (*SCN1A, SCN2A, SCN1B, GABRA1, GABRG2, HCN1, CHD2* and *PCDH19*). We identified two *de novo* variants in *SCN1A* in two patients (Figure 3).

**Figure 3:**
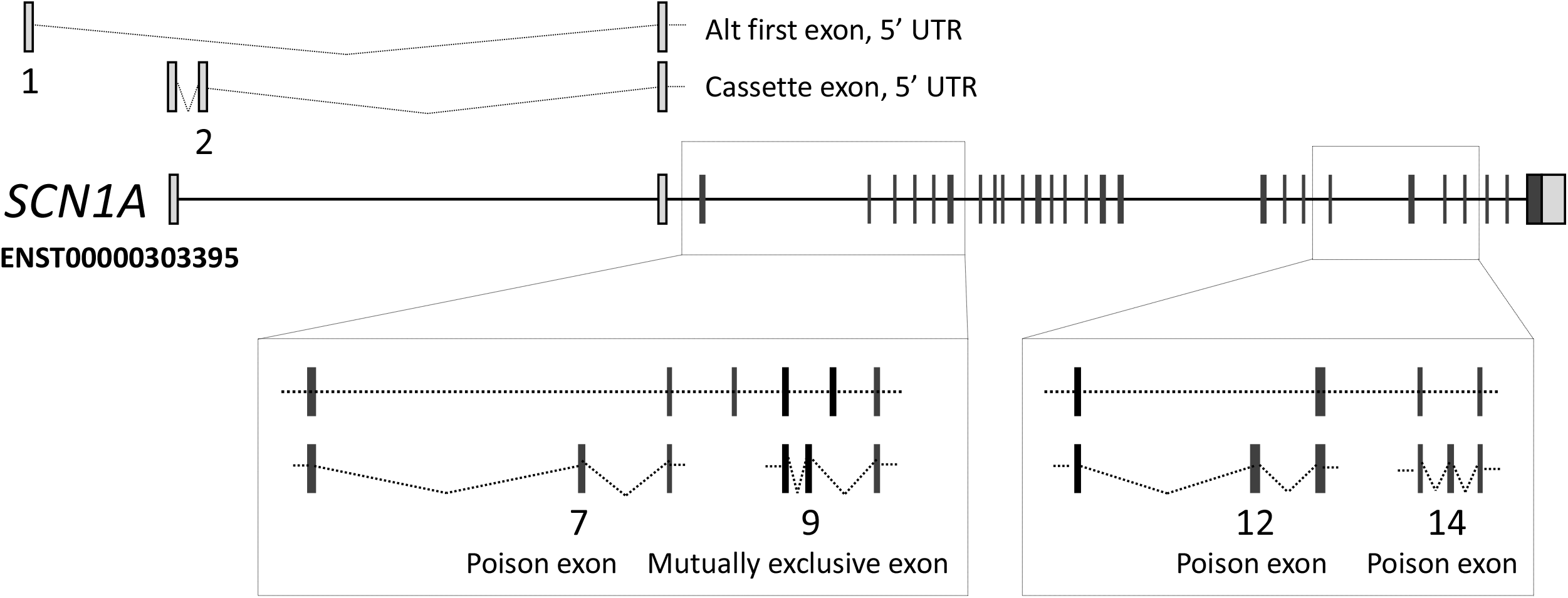
Variants in novel coding regions are associated with Dravet syndrome. (a) Pedigrees and Sanger sequencing traces of the two families with a *de novo SCN1A* variant in the novel identified poison exons. (b) The two transcripts containing the variants, relative to the full-length transcript. Red exons are coding, white exons are non-coding. (c) Variants are predicted to disrupt a hnRNP A1 recognition site.

Patient 1 is a fifteen-year-old girl diagnosed with Dravet syndrome. Previous screening efforts did not reveal a molecular diagnosis (including *SCN1A, STXBP1* and a gene panel^30^ consisting of known and candidate genes for Dravet syndrome and Myoclonic Atonic Epilepsy). The *de novo* variant identified here is found in an *SCN1A* poison exon (feature 7 originally in intron 1; GRCh38 chr2: 166060831, ENST00000636759.1:c.301G>A; Table 2, Figure 2, Figure 3). The variant was validated with Sanger sequencing, and maternity and paternity was confirmed using an in-house developed multiplex PCR panel consisting of 16 STR-markers scattered over the genome, including the X and Y chromosome. She is the only child of healthy non-consanguineous parents. The father had photosensitive epilepsy as a child but is now seizure-free without medication. A half-sister on the mother’s side has epilepsy but no developmental delay. Due to the normal development of both the father and the maternal half-sister, and due to the lack of kinship between father and half-sister, it is not expected that the family members share the same genetic variant as the proband to explain their epilepsy. However, it cannot be excluded that the father has a low-grade mosaicism for the *SCN1A* variant that was not detected through Sanger sequencing. The proband first presented with febrile seizures when she was 8 months old. She had focal impaired awareness seizures and later also developed afebrile tonic-clonic seizures starting at 18 months old that occurred very frequently until the age of 5 years. She also had absences and myoclonic seizures. Electroencephalograms (EEG) showed background slowing and paroxysmal slow spike and spike waves. Development was normal prior to seizure onset, but slowed soon after, resulting in moderate intellectual disability. Brain MRI imaging showed no abnormalities and normal spectroscopic sequences. She is currently being treated with a combination of levetiracetam, topiramate, clobazam and stiripentol, but still has frequent tonic-clonic seizures.

Patient 2 is a fourteen-year-old girl, diagnosed with Dravet syndrome. Previous screening of *SCN1A, PCDH19* and *HCN1* did not result in the identification of a pathogenic variant. Our study identified a *de novo* variant in a different poison exon of *SCN1A* (feature 14 originally in intron 22; GRCh38 chr2: 165999107, ENST00000635893.1:c.79G>A; Table 2, Figure 2, Figure 3). The variant was validated with Sanger sequencing, and maternity and paternity was confirmed using the same in-house developed multiplex PCR panel. She is born from healthy non-consanguineous parents. There is no familial history of epilepsy. The proband had her first febrile seizure evolving to status epilepticus when she was six months old. She further had on average six or seven episodes of generalized tonic-clonic status epilepticus per year, mostly with fever. After seizure onset her development slowed, resulting in moderate intellectual disability. Other comorbidities include ataxia, orofacial dyspraxia, difficulties with fine motor skills, hyperkinesia and sleep disturbances. EEG at the age of 5 years showed bifrontal slow waves and rare temporal spikes. MRI was normal apart from an asymptomatic pituitary cyst. She is currently being treated with a combination of valproate, clobazam and topiramate, which has reduced seizure frequency.

To quantify the significance of *de novo* mutation enrichment in the novel 450bp of CDS sequences (sum of features 7, 9, 12, and 14 from Table 2) and to estimate the probability of identifying 2 or more *de novo* variants in a cohort of 122 sequenced probands, we performed a *de novo* mutation enrichment statistical test using the fitDNM package^31^. This package returns the Poisson unweighted P-value based on expected mutability rate and shows that it is improbable to observe 2 *de novo* variants among 122 individuals along this stretch of 450bp of CDS (unadjusted p = 9.5×10^-7^). Even after conservative exome-wide multiple testing correction for 18,000 possible protein-coding genes, this remains significant (p = 0.017), confirming a significant enrichment of *de novo* variants in the novel *SCN1A* sequences.

Neither poison exon is well-conserved (PhastCons scores 0.102 and 0.000 for the two variants respectively). Both variants were predicted to alter binding sites for hnRNP A1, which is a splice ‘silencer’ that promotes exon skipping^32,33^. We therefore postulate that the alterations of these respective motifs could lead to increased inclusion of the poison exons, and therefore a reduction in the production of functional *SCN1A* protein due to NMD. Further functional work will be needed, however, to validate this hypothesis.

## Discussion

The ability to infer clinical information about a patient’s genome relies upon reference data sets that help to make informed decisions on putative causative variants. Accordingly, the confidence of such decisions is dependent upon the reliability of the underlying data, against which a patient’s genome is analysed. For example, the human genome reference sequence is still incomplete, which is demonstrated by the ongoing work that the Genome Reference Consortium is undertaking to fill remaining sequence gaps in the human genome, as well as representing different population haplotypes^9^. However, the incompleteness of the human transcriptome should also be a consideration for genome interpretation, since a complete set of transcripts from all the different tissue types and developmental stages that are naturally present in the human genome is not yet available. Until it is possible to confidently generate a *de novo* genome sequence for a patient, in conjunction with a complete transcriptome (and also proteome etc), researchers and clinicians must rely upon data that is available in public databases.

As such, our investigation shows that the current set of human transcripts, while being expertly curated and maintained, cannot be considered complete. To illustrate this, we have used human brain-transcriptomic data to rigorously interrogate the human reference transcriptome for missing transcriptional features. Interestingly, the extended transcriptional footprint of these genes immediately allowed for 294 intronic or intergenic variants, found in human mutation databases, to be reclassified as exonic, while a further 70 intronic variants were reclassified as splice-site proximal.

To further our study, we investigated a cohort of people with Dravet syndrome, which has a robust phenotype/genotype correlation with *SCN1A* using our updated transcriptome data; the presumption being that any new exon features that capture a variant in *SCN1A* could support a diagnosis. On investigation, we identified three variants, within three distinct poison exons of the *SCN1A* gene, that are associated with Dravet syndrome, each of which is absent from current diagnostic tests. Variants in poison exons have been previously described to cause Mendelian disorders, *via* mechanisms proposed to affect the level of protein expression^34,35^. Haploinsufficiency of *SCN1A* is a cause of Dravet syndrome, hence it seems reasonable to speculate that a higher inclusion of any of these poison exons could lead to a net reduction of functional protein and thus the disease phenotype; this has in fact now been established for ClinVar variant RCV000209951 in the poison exon described by Carvill et al^29^. For the two *de novo* variants first reported here, we recognise that similar efforts to ascribe true pathogenicity will now be required, which is complicated by the neuron-specific expression pattern of *SCN1A*. However, given the strong and clearly established genotype-phenotype correlation between Dravet syndrome and *SCN1A*, it is appropriate to consider these two variants as strong candidates for driving pathogenicity, not least because making this genetic diagnosis can underpin potentially life-saving changes to medication, as well as informing prognosis and stopping further unnecessary investigations. We emphasise that these are *de novo* variants, and their absence in approximately 15,500 genomes in gnomAD^36^ is consistent with negative selection in human populations. Also, the observation of multiple disease-linked variants within the total 450bp space of predicted novel coding, or NMD-triggering sequence, is highly improbable among just 122 probands -considering also that the average human is expected to have fewer than 100 single nucleotide *de novo* variants in total -which suggests they have been identified due to the focus on a cohort of molecularly undiagnosed Dravet Syndrome probands.

Nonetheless, it is striking that only the poison exon overlapping ClinVar variant RCV000209951 exhibits notable evolutionary conservation. This point is of immediate practical importance because the *de novo* variants reported here are at risk of being filtered out by prioritisation algorithms that utilise conservation metrics. These findings may also be surprising from a biological point of view, i.e. considering potential mechanisms of pathogenicity, given the traditional weight placed on the maxim that ‘conservation = function’. However, while gene-level conservation is typically studied in the context of protein-coding sequences, far less is known about the evolutionary dynamics of gene regulatory programs linked to alternative splicing. These concepts may be especially relevant to the brain, which is known to be particularly rich in alternative splicing compared to other organs and tissues. Indeed, given that the human brain has evolved and enlarged considerably with respect to apes over the past 2.5 million years^36^, we can speculate that novel functionality is generally linked to newly evolved sequences^37^. Nonetheless, we also observed that the transcriptional support for both poison exons is not strong. These observations would be reconciled with our inferences into pathogenicity if the *de novo* mutations do indeed turn out to lead to increased inclusion of the poison exons, and it would therefore be informative to know whether these exons have higher levels of inclusion in the two Dravet patients.

Extrapolating from this discussion, it is reasonable to consider that all of the 3,550 transcripts annotated here, within an exemplar set of 191 genes associated with epilepsy, are of *potential* clinical interest. However, we recognise that detailed work will be required to establish which of these variants have a true clinical association with epilepsy, as well as the biological mechanisms by which they drive disease. A logical next step would be to resequence these novel regions in patients with epilepsy that currently lack a molecular diagnosis.

Furthermore, we recognise that the value of adding large numbers of additional transcribed regions to disease-linked genes could be questioned while it remains unclear exactly which transcript models have biological relevance. Broadly speaking, diagnostics based on expanded gene annotation has the potential to reduce false negative variant interpretations (i.e. to incorporate important ‘missed’ variants), but at the expense of an increase in false positives. As discussed, our *SCN1A* variant interpretations benefit from the high concordance between perturbations to this gene and Dravet syndrome, as well as the fact that these are *de novo* mutations. It is less certain how expanded annotation would perform in clinical scenarios where this is not the case. For example, hundreds of genes have now been linked to autism spectrum disorder with varying degrees of confidence and this disorder has a far more heterogeneous causation than Dravet syndrome. In our view, identified sequences with strong markers for functionality, especially those that can be established as coding sequences based on evolutionary conservation and/or proteomics data, should be considered those most likely to have functionality and therefore those with the most potential value in the search for undiscovered disease-linked variants. Nonetheless, our work here illustrates that a consideration of the full transcriptional profile of a gene can also be fruitful, i.e. including transcripts with poor conservation and weak transcription, and those that do not appear to encode proteins. As discussed, we believe this point is particularly apposite when considering that pathogenicity can also arise from gain of function modes.

Finally, establishing a genetic cause for epilepsy, in an individual, is a key step in clinical management. It provides an explanation, terminates the diagnostic odyssey and may inform treatment options and prognostication. Furthermore, it helps with determining other management issues (e.g. known associated comorbidities like gait disorders, involvement of other organs, dysphagia), genetic counselling and overall care, where having a name for a condition typically facilitates access to services. Last, but not least, it provides a label and typically relieves parental guilt. Current methods of establishing a genetic diagnosis in clinical settings consist of candidate gene analysis, gene panel or WES and array comparative genomic hybridization to identify copy number variants. If these methods are undertaken and no variant is found, such cases usually go into research projects, which generally take longer and are more uncertain. If the original tests to establish a diagnosis are missing annotation, this all leads to costly and unnecessary delays, both for patients, their family and for the healthcare system. This study further raises the question whether it would be more desirable to use WGS for diagnostic purposes, as data can be iteratively re-annotated when updated annotation information becomes available. Similarly, this approach has already been proven successful as iteratively re-analyzing patient genomes when new causative genes are discovered increases diagnostic yield^6,38^.

In summary, our findings suggest that there are potentially additional causative genetic variants to be identified in and around epilepsy-related genes such as *SCN1A*, including in predicted poison exons. We anticipate that the inclusion of the novel transcripts identified here will further increase the number of variants found in *SCN1A* in people with Dravet Syndrome. Furthermore, if the same approach we have taken to *SCN1A* and Dravet is applied to other focussed cohorts for the other 190 genes in our study, we would expect to find newly explained cases in the gene that was originally suspected by the clinician.

## Supporting information

Supplemental gene updates

Supplemental statistical analysis

## Acknowledgments

We thank the following: all patients and their families who participated in this study, as well as the teams who were involved in recruiting patients and gathering samples and data at the respective study sites; Dr Gautam Ambegaonkar, Ms Margo Elsworth, Dr Jill Gordon, Dr Alasdair Parker, Dr Elizabeth Radford, Ms Kuldeep Stohr (NHS England) for clinical support and advice; Dr Andrew Jaffe (Lieber Institute for Brain Development, USA) for advice with utilising the 6 different life-stage transcriptomic datasets used in this study; Dr Matthew Hurles (Wellcome Sanger Institute, UK) for initial discussions regarding this study. We also thank Imogen and Jasper Steward, both of whom have rare neurological disorders and have been instrumental in driving this study.

## Informed Consent

All the patients/parents in this study have given informed consent. Sequencing was done after ethical approval from the ERC of the University of Antwerp

## Author Contributions

Conceiving the project: CAS, DK, JH, BAM, DG, FLR, SMS, AF. Planning and designing the study: CAS, JR, JMM, JH, BAM, DG, FLR, NJL, SW, PDJ, SMS, AF. Gene annotation: CAS, MMS, JMM. Data analysis: JMG, BUR, DP, SF, ET, ASJ, ASR, JW, JC, MD, RG, RP, DV, SP. Patient DNA collection and clinical examination: HS, BC, PL, CN, AL, SW, PDJ. Wet-lab experiments and analysis of patient DNA: JR, MV. Writing manuscript: CAS, JR, JMG, BUR, DP, JW, JC, MD, RG, RP, SP, JMM, BAM, FLR, NJL, SW, PDJ, SMS, AF. Contributed resources: SA, NJL. Drafting, reviewing and approval of the manuscript: all authors.

## Online methods

### Selection of genes

191 genes associated with deleterious variants implicated in epilepsy in general and EIEE, in particular, were included in this study. These included sixty-six genes from the Great Ormond Street Hospital (GOSH) for Children, London, UK, EIEE gene-panel (http://www.labs.gosh.nhs.uk/media/759010/eiee_v7.pdf), selected as established causes of early-onset seizures and/or severe developmental delay in patients without frequent major structural brain anomalies. Genes leading to neurometabolic disorders with readily identifiable blood/urine/cerebrospinal fluid (CSF) biomarkers were not included. An additional fifty eight genes were included from the Addenbrooke’s Hospital, Cambridge, UK, EIEE gene panel, as well as 14 genes from the DDD study^39^ and 53 genes from literature searching. The full list can be found in the Supplemental Data 2.

### Gene annotation

Manual re-annotation of the 191 genes was performed on GRCh38 (https://www.ncbi.nlm.nih.gov/grc/human) according to the guidelines of the HAVANA (Human And Vertebrate Analysis and Annotation) group^40^ (and ftp://ftp.sanger.ac.uk/pub/project/havana/Guidelines/Guidelines_March_2016.pdf). In summary, the HAVANA group produces annotation, largely based on the alignment of transcriptomic (ESTs and mRNAs) and protein sequence data from GenBank^41^ and Uniprot^42^. These data were aligned to the individual BAC clones that make up the reference genome sequence using BLAST^43^ with a subsequent realignment of transcript data by Est2Genome^44^. In addition, SLR-RNA-Seq data^45^ mapped using Gmap^46^, PACBIO^47^ reads mapped using STAR^48^, and fetal and infant RNA-Seq data^17^ mapped using cufflinks^49^ were also used to identify novel transcripts and splice junctions. Data are available at www.gencodegenes.org/releases/. Updated annotation of the 191 genes described in this study on GRCh38 is represented in GENCODE releases from v27 (August 2017) onwards. In addition, the updated annotation is available remapped to GRCh37 [https://github.com/diekhans/gencode-backmap] here: http://www.gencodegenes.org/releases/grch37_mapped_releases.html.

### Identification of novel splicing events

Transcript structures in public releases of GENCODE before (GENCODE v20) and after (GENCODE v28) the manually updated annotations were compared to find novel exons, novel introns and shifted splice junctions. The number of genomic bases covered by the extended gene annotation and novel coding sequence was calculated using a custom Perl script. Novel exons are defined as those in the updated annotation that shared no sequence with any exon in the old annotation. Novel introns were those introns in the updated annotation that did not match exactly an intron in the old annotation. Shifted splice junctions occurred when an exon in the updated annotation shares an overlap with an exon in the old annotation, but at least one of the splice sites was not in the same location. Novel retained intron transcripts were excluded from this analysis. RefSeq annotation for transcript counting was extracted from the Ensembl release 84 (March 2016) “RefSeq GFF3 annotation import”.

### Identification of novel transcriptional features on GRCh38 using Fetal and Infant RNA-Seq data

Illumina data from Jaffe et al^17^ was re-mapped for fetal and infant transcriptome to GRCh38 to identify novel transcriptional features. Fastq files from the following datasets were downloaded from ENA: SRR1554537, SRR1554538, SRR1554541, SRR1554544, SRR1554546, SRR1554549, SRR1554551, SRR1554553, SRR1554554, SRR1554566, SRR1554567 and SRR1554568. Data were mapped to GRCh38 with TopHat (tophat-2.0.13)^50^. Reads mapping to the gene regions that were studied, were merged into two files containing fetal and infant alignments using SAMtools^51^. Transcript models were generated from the fetal and infant BAM files using Cufflinks (cufflinks-2.2.1)^50^. Novel introns and splice sites were identified, a BED file generated and passed to manual annotators for checking.

### Quantification of gene expression at exon level

Raw reads from Jaffe et al^17^ were available *via* study accession SRP045638. The 36 paired-end libraries were analyzed using the iRAP pipeline (https://github.com/nunofonseca/irap). First, raw reads in the original FASTQ files underwent quality assessment and filtering^52^. They were then aligned against the GRCh38 genome reference using Tophat2^53^, with the option: ‘--min-intron-length 6’.

### Analysis of expression of novel splice features

Integrative Pipeline for Splicing Analyses (IPSA)^54^ was employed to produce splice junctions and their read counts from Tophat2 alignments of Jaffe et al^17^ on GRCh38 human genome. This analysis included 36 human brain pre-frontal cortex samples, corresponding to six different developmental stages (Fetal, Infant, Child, Teen, Adult, Old)^17^. IPSA was run with the default parameters and the pre-annotation update release GENCODE v20 as a reference. Transcript expression levels around introns were estimated from the number of reads supporting the respective splice junction. Expression of splice junctions was normalized by the total number of reads in each sample.

### Mapping annotation from reference human genome to mouse genome

The TransMap cross-species alignment algorithm was used to map all annotated transcripts from the reference human genome (GRCh38) to the reference mouse genome (GRCm38). The alignments are created using synteny-filtered pairwise genome alignments (chains and nets) produced using BLASTZ^21,55,56^. All transcript models mapped to mouse were manually-assessed to identify failures to align correctly at the base, exon and intron level.

### Identification of conservation of novel coding sequence

The novel EIEE gene annotation was obtained by subtracting the GENCODE v20 gene annotation from the current one (equivalent to GENCODE v28) using “bedtools subtract” separately for exon and CDS regions. The RepeatMasker repeat features (except low complexity elements) extracted from Ensembl were subsequently subtracted from this novel annotation. The filtered novel annotation was then intersected with:

a. phastConsElements100way.bed obtained from the UCSC Table Browser (https://genome.ucsc.edu/cgi-bin/hgTables);
b. 27 amniota vertebrates GERP constrained elements from Ensembl (ftp://ftp.ensembl.org/pub/release-90/bed/ensembl-compara/27_amniota_vertebrates_gerp_constrained_elements/gerp_constrained_elements.homo_sapiens.bed.gz);
c. PhyloCSF (58 mammals) approximate coding regions from the PhyloCSF track hub (https://data.broadinstitute.org/compbio1/PhyloCSFtracks/trackHub/hg38/trackDb.txt). In all cases the overlap with the novel EIEE gene annotation was carried out using “bedtools intersect”.

### Identification of variants in updated annotation

The collection of variants available under “All phenotype-associated -short variants (SNPs and indels)” in Ensembl release 90 (August 2017) was intersected separately with the EIEE-related gene exons in GENCODE 20 and in the current annotation, using a custom Perl script and the Ensembl API. The variants overlapping the current annotation but not GENCODE 20 were reported. A second round was carried out considering only the 8-nt exon flanking sequences in both annotation sets.

As a proof-of-concept, we screened a cohort of 122 patients with Dravet syndrome or a clinically similar severe myoclonic epilepsy phenotype, for the 125 novel regions identified from this study of all genes that have previously been associated with Dravet syndrome (*SCN1A, SCN2A, SCN1B, GABRA1, GABRG2, HCN1, CHD2* and *PCDH19*), using an amplicon targeted amplification assay (Agilent, https://www.agilent.com). All samples underwent diagnostic screening for *SCN1A* (including both sequencing and CNV analysis), but no pathogenic variants had been identified, after which they were included in research. Additionally, several patients were screened for genetic variants in epilepsy-associated genes using Sanger sequencing, gene panels or WES.

Primers for the multi-amplicon target panel were designed using the mPCR software (Agilent)^57^. Specific target regions were amplified using multiplex PCR, followed by purification of the equimolar pooled amplicons using Agencourt AMPureXP beads (Beckman Coulter, CA, USA). Individual barcodes (Illumina Nextera XT) were incorporated in a universal PCR step prior to sample pooling. Libraries were sequenced on a MiSeq platform using v3 reagent kit with a paired-end read length of 300 bp (Illumina, USA).

Analysis was performed in-house with a standardized pipeline integrated in genomecomb^58^. The pipeline used fastq-mcf (http://code.google.com/p/ea-utils) for adapter clipping. Reads were then aligned using bwa mem^59^ and the resulting sam file converted to bam using Samtools^51^. Bam files were sorted using biobambam^60^. Realignment in the neighbourhood of indels was performed with GATK^61^. Amplicon primers were clipped using genomecomb^58^. Variants were called at all positions with a total coverage >=5 using both GATK^61^ and Samtools^51^. At this initial stage, positions with a coverage <5 or a score <30 were considered unsequenced. The resulting variant sets of different individuals were combined and annotated and filtered using genomecomb^58^.

Variants with a coverage above 7 and a GATK quality score above 50, that were identified by both variant callers (GATK and Samtools) and were absent in publicly available databases (EXaC^62^, gnomAD^62^, 1000 Genomes^63^, Exome Variant Server (http://evs.gs.washington.edu/EVS/)) and our in house database, were extracted. Furthermore, variants present in homopolymer regions (>8 homopolymers) or simple repeat regions, were excluded. The effect of the variant was predicted with the Ensembl Variant Effect Predictor (VEP)^64^, using the manually annotated GENCODE dataset as custom gene annotation. Variants were validated and the segregation was checked using bi-directional Sanger sequencing. For *de novo* variants, maternity and paternity was confirmed using an in-house developed multiplex PCR panel consisting of 16 STR-markers scattered over the genome, including the X and Y chromosome.

To predict the effect of the *de novo* variants on splicing efficiency, we used the quick mutation analysis from human splice finder^65^, using the CDS from the NMD transcripts as custom sequence input.

### Testing for significant excess of *de novo* mutation among novel *SCN1A* CDS protein-coding sequence

The genomics sequence corresponding to novel protein-coding features of *SCN1A* (Features 7, 9, 12 and 14; Table 2) were concatenated to reflect a consecutive test sequence of length 450bp on hg38. To test whether the observed number of 2 *de novo* mutations among this stretch of 450bp was significant, based on a sampled cohort of 122 individuals, we adopted a modified version of R package fitDNM^31^. The published fitDNM provides both a PolyPhen-2 weighted and Poisson unweighted p-value. Here, since PolyPhen-2 scores do not exist for the entirety of the novel CDS sequence we focus on the Poisson unweighted P-value. The modification to the original fitDNM package was to correct a type conversion error, which possibly occurred due to different versions of R used for our analysis (v3.5.0) compared to the original package version. Specifically, we fixed a clash where a ‘T’-valued variable (intended for Thymine) was handled as “TRUE”. The adapted fitDNM package accompanied by input and output files from this analysis are available upon request. This R package then takes as input the underlying mutability of the 450bp (Supplemental Data 3), the total number of observed *de novo* mutations among that 450 bases of sequence and the total number of probands tested for a *de novo* mutation in that 450 bases of sequence. Subsequently, we conservatively corrected the resulting Poisson unweighted P-value by 18,000 to reflect approximately the total number of WES studied protein-coding genes in the human exome.

## References

1. Nieh, S.E. & Sherr, E.H. Epileptic encephalopathies: new genes and new pathways. Neurotherapeutics 11, 796–806 (2014).

2. EuroEpinomics-R.E.S.Consortium, EpilepsyPhenome/Genome-Project & Epi4K. Consortium. De novo mutations in synaptic transmission genes including DNM1 cause epileptic encephalopathies. Am J Hum Genet 95, 360–70 (2014).

3. Deciphering-Developmental-Disorders-Study. Prevalence and architecture of de novo mutations in developmental disorders. Nature 542, 433–438 (2017).

4. Mark, C. et al. The 100,000 Genomes Project Protocol, (2017).

5. EpiPMConsortium. A roadmap for precision medicine in the epilepsies. Lancet Neurol 14, 1219–28 (2015).

6. Wright, C.F. et al. Making new genetic diagnoses with old data: iterative reanalysis and reporting from genome-wide data in 1,133 families with developmental disorders. Genet Med (2018).

7. Frankish, A. et al. GENCODE reference annotation for the human and mouse genomes. Nucleic Acids Res 47, D766–D773 (2019).

8. O’Leary, N.A. et al. Reference sequence (RefSeq) database at NCBI: current status, taxonomic expansion, and functional annotation. Nucleic Acids Res 44, D733–45 (2016).

9. Schneider, V.A. et al. Evaluation of GRCh38 and de novo haploid genome assemblies demonstrates the enduring quality of the reference assembly. Genome Res 27, 849–864 (2017).

10. Imanishi, T. et al. Integrative annotation of 21,037 human genes validated by fulllength cDNA clones. PLoS Biol 2, e162 (2004).

11. Batzoglou, S., Pachter, L., Mesirov, J.P., Berger, B. & Lander, E.S. Human and mouse gene structure: comparative analysis and application to exon prediction. Genome Res 10, 950–8 (2000).

12. Reggiani, C. et al. Novel promoters and coding first exons in DLG2 linked to developmental disorders and intellectual disability. Genome Med 9, 67 (2017).

13. Epilepsy-Genetics-Initiative. De novo variants in the alternative exon 5 of SCN8A cause epileptic encephalopathy. Genet Med 20, 275–281 (2018).

14. Mudge, J.M. et al. The origins, evolution, and functional potential of alternative splicing in vertebrates. Mol Biol Evol 28, 2949–59 (2011).

15. Kurosaki, T., Popp, M.W. & Maquat, L.E. Quality and quantity control of gene expression by nonsense-mediated mRNA decay. Nat Rev Mol Cell Biol (2019).

16. Anna, A. & Monika, G. Splicing mutations in human genetic disorders: examples, detection, and confirmation. J Appl Genet 59, 253–268 (2018).

17. Jaffe, A.E. et al. Developmental regulation of human cortex transcription and its clinical relevance at single base resolution. Nat Neurosci 18, 154–161 (2015).

18. Mercer, T.R., Dinger, M.E., Sunkin, S.M., Mehler, M.F. & Mattick, J.S. Specific expression of long noncoding RNAs in the mouse brain. Proc Natl Acad Sci U S A 105, 716–21 (2008).

19. Young, R.S. & Ponting, C.P. Identification and function of long non-coding RNAs. Essays Biochem 54, 113–26 (2013).

20. Frankish, A., Mudge, J.M., Thomas, M. & Harrow, J. The importance of identifying alternative splicing in vertebrate genome annotation. Database (Oxford) 2012, bas014 (2012).

21. Stanke, M., Diekhans, M., Baertsch, R. & Haussler, D. Using native and syntenically mapped cDNA alignments to improve de novo gene finding. Bioinformatics 24, 637–44 (2008).

22. Siepel, A. et al. Evolutionarily conserved elements in vertebrate, insect, worm, and yeast genomes. Genome Res 15, 1034–50 (2005).

23. Lin, M.F., Jungreis, I. & Kellis, M. PhyloCSF: a comparative genomics method to distinguish protein coding and non-coding regions. Bioinformatics 27, i275–82 (2011).

24. Dravet, C. The core Dravet syndrome phenotype. Epilepsia 52 Suppl 2, 3–9 (2011).

25. Djemie, T. et al. Pitfalls in genetic testing: the story of missed SCN1A mutations. Mol Genet Genomic Med 4, 457–64 (2016).

26. Parihar, R. & Ganesh, S. The SCN1A gene variants and epileptic encephalopathies. J Hum Genet 58, 573–80 (2013).

27. Tate, S.K. et al. Genetic predictors of the maximum doses patients receive during clinical use of the anti-epileptic drugs carbamazepine and phenytoin. Proc Natl Acad Sci U S A 102, 5507–12 (2005).

28. Oh, Y. & Waxman, S.G. Novel splice variants of the voltage-sensitive sodium channel alpha subunit. Neuroreport 9, 1267–72 (1998).

29. Carvill, G.L. et al. Aberrant Inclusion of a Poison Exon Causes Dravet Syndrome and Related SCN1A-Associated Genetic Epilepsies. Am J Hum Genet 103, 1022–1029 (2018).

30. Carvill, G.L. et al. Targeted resequencing in epileptic encephalopathies identifies de novo mutations in CHD2 and SYNGAP1. Nat Genet 45, 825–30 (2013).

31. Jiang, Y. et al. Incorporating Functional Information in Tests of Excess De Novo Mutational Load. Am J Hum Genet 97, 272–83 (2015).

32. Jean-Philippe, J., Paz, S. & Caputi, M. hnRNP A1: the Swiss army knife of gene expression. Int J Mol Sci 14, 18999–9024 (2013).

33. Beusch, I., Barraud, P., Moursy, A., Clery, A. & Allain, F.H. Tandem hnRNP A1 RNA recognition motifs act in concert to repress the splicing of survival motor neuron exon 7. Elife 6(2017).

34. Zhang, X. et al. Cell-Type-Specific Alternative Splicing Governs Cell Fate in the Developing Cerebral Cortex. Cell 166, 1147–1162 e15 (2016).

35. Lynch, D.C. et al. Disrupted auto-regulation of the spliceosomal gene SNRPB causes cerebro-costo-mandibular syndrome. Nat Commun 5, 4483 (2014).

36. Hofman, M.A. Evolution of the human brain: when bigger is better. Front Neuroanat 8, 15 (2014).

37. Levchenko, A., Kanapin, A., Samsonova, A. & Gainetdinov, R.R. Human Accelerated Regions and Other Human-Specific Sequence Variations in the Context of Evolution and Their Relevance for Brain Development. Genome Biol Evol 10, 166–188 (2018).

38. Costain, G. et al. Periodic reanalysis of whole-genome sequencing data enhances the diagnostic advantage over standard clinical genetic testing. Eur J Hum Genet (2018).

39. Wright, C.F. et al. Genetic diagnosis of developmental disorders in the DDD study: a scalable analysis of genome-wide research data. Lancet 385, 1305–14 (2015).

40. Harrow, J. et al. GENCODE: the reference human genome annotation for The ENCODE Project. Genome Res 22, 1760–74 (2012).

41. Benson, D.A. et al. GenBank. Nucleic Acids Res 45, D37–D42 (2017).

42. Gaulton, A. et al. The ChEMBL database in 2017. Nucleic Acids Res 45, D945–D954 (2017).

43. Altschul, S.F. et al. Gapped BLAST and PSI-BLAST: a new generation of protein database search programs. Nucleic Acids Res 25, 3389–402 (1997).

44. Mott, R. EST_GENOME: a program to align spliced DNA sequences to unspliced genomic DNA. Comput Appl Biosci 13, 477–8 (1997).

45. Tilgner, H. et al. Comprehensive transcriptome analysis using synthetic long-read sequencing reveals molecular co-association of distant splicing events. Nat Biotechnol 33, 736–42 (2015).

46. Wu, T.D., Reeder, J., Lawrence, M., Becker, G. & Brauer, M.J. GMAP and GSNAP for Genomic Sequence Alignment: Enhancements to Speed, Accuracy, and Functionality. Methods Mol Biol 1418, 283–334 (2016).

47. Rhoads, A. & Au, K.F. PacBio Sequencing and Its Applications. Genomics Proteomics Bioinformatics 13, 278–89 (2015).

48. Dobin, A. et al. STAR: ultrafast universal RNA-seq aligner. Bioinformatics 29, 15–21 (2013).

49. Trapnell, C. et al. Transcript assembly and quantification by RNA-Seq reveals unannotated transcripts and isoform switching during cell differentiation. Nat Biotechnol 28, 511–5 (2010).

50. Trapnell, C. et al. Differential gene and transcript expression analysis of RNA-seq experiments with TopHat and Cufflinks. Nat Protoc 7, 562–78 (2012).

51. Li, H. et al. The Sequence Alignment/Map format and SAMtools. Bioinformatics 25, 2078–9 (2009).

52. Petryszak, R. et al. Expression Atlas update--a database of gene and transcript expression from microarray-and sequencing-based functional genomics experiments. Nucleic Acids Res 42, D926–32 (2014).

53. Trapnell, C., Pachter, L. & Salzberg, S.L. TopHat: discovering splice junctions with RNA-Seq. Bioinformatics 25, 1105–11 (2009).

54. Pervouchine, D.D., Knowles, D.G. & Guigo, R. Intron-centric estimation of alternative splicing from RNA-seq data. Bioinformatics 29, 273–4 (2013).

55. Zhu, J. et al. Comparative genomics search for losses of long-established genes on the human lineage. PLoS Comput Biol 3, e247 (2007).

56. Siepel, A. et al. Targeted discovery of novel human exons by comparative genomics. Genome Res 17, 1763–73 (2007).

57. Goossens, D. et al. Simultaneous mutation and copy number variation (CNV) detection by multiplex PCR-based GS-FLX sequencing. Hum Mutat 30, 472–6 (2009).

58. Reumers, J. et al. Optimized filtering reduces the error rate in detecting genomic variants by short-read sequencing. Nat Biotechnol 30, 61–8 (2012).

59. Li, H. & Durbin, R. Fast and accurate short read alignment with Burrows-Wheeler transform. Bioinformatics 25, 1754–60 (2009).

60. Tischler, G. & Leonard, S. biobambam: tools for read pair collation based algorithms on BAM files. Source Code for Biology and Medicine 9, 13 (2014).

61. DePristo, M.A. et al. A framework for variation discovery and genotyping using nextgeneration DNA sequencing data. Nat Genet 43, 491–8 (2011).

62. Lek, M. et al. Analysis of protein-coding genetic variation in 60,706 humans. Nature 536, 285–91 (2016).

63. Genomes Project, C. et al. A global reference for human genetic variation. Nature 526, 68–74 (2015).

64. McLaren, W. et al. The Ensembl Variant Effect Predictor. Genome Biol 17, 122 (2016).

65. Desmet, F.O. et al. Human Splicing Finder: an online bioinformatics tool to predict splicing signals. Nucleic Acids Res 37, e67 (2009).

