## Supplementary figures and images for "Systematic re-annotation of 191 genes associated with early-onset epilepsy unmasks *de novo* variants linked to Dravet syndrome in novel *SCN1A* exons"

### Supplemental Figure 1

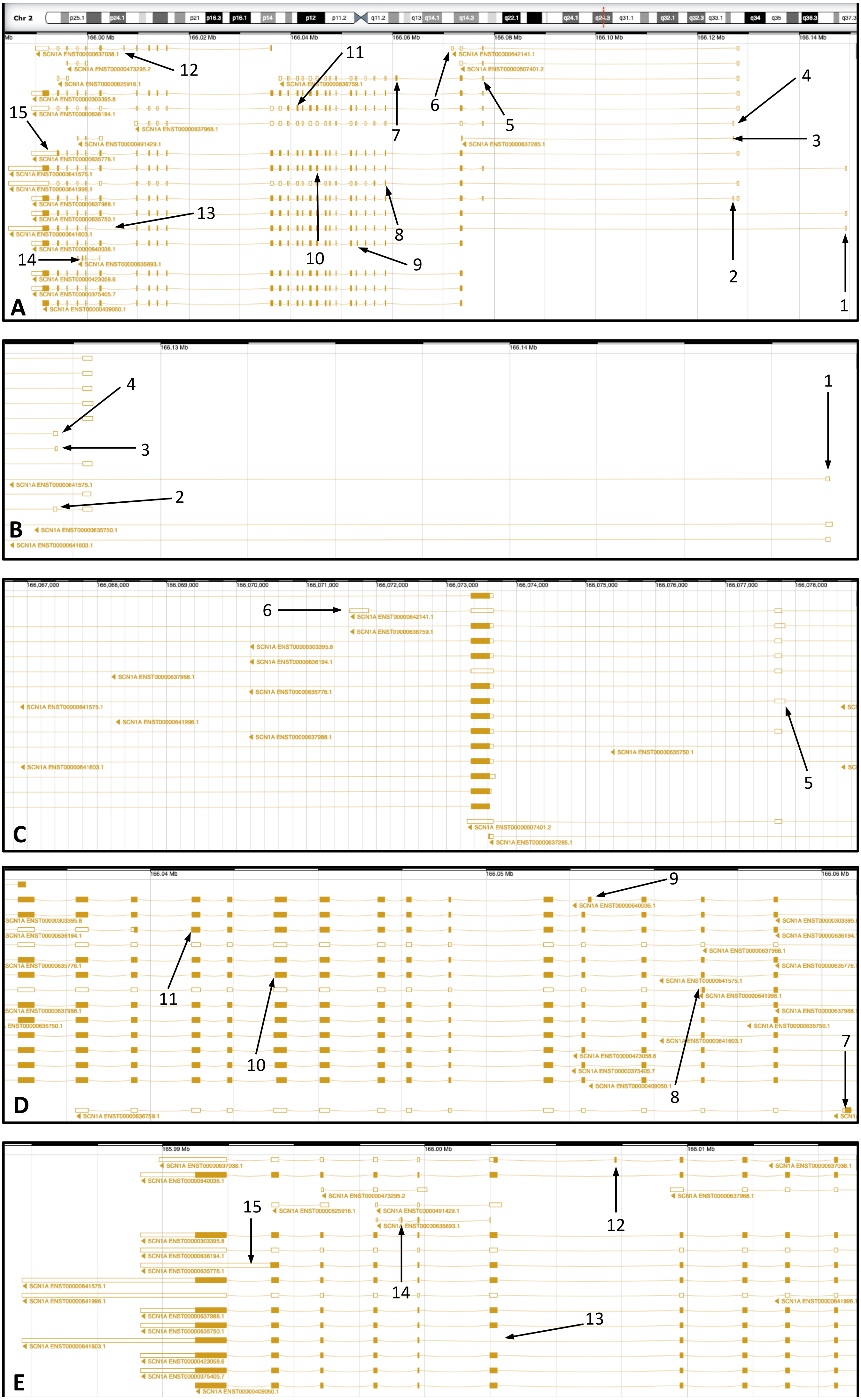
